# Dietary and Biological Assessment of Omega-3 Status of Collegiate Athletes: A Cross-Sectional Analysis

**DOI:** 10.1101/2020.01.27.920991

**Authors:** Peter P. Ritz, Mark B. Rogers, Jennifer S. Zabinsky, Valisa E. Hedrick, John A. Rockwell, Ernest Rimer, Samantha Kostelnik, Matthew W. Hulver, Michelle S. Rockwell

## Abstract

Omega-3 fatty acids (ω-3 FA) play a number of important functions in health and human performance. While previous research has suggested that low ω-3 FA status is prevalent in the general population, little information about athletes’ ω-3 FA status is available. The purpose of this study was to assess the omega-3 fatty acid (ω-3 FA) status of collegiate athletes. Dietary ω-3 FA intake was evaluated in athletes from nine NCAA Division I institutions (n=1,528, 51% male, 19.9 ± 1.4 years of age, 29 sports represented) via food frequency questionnaire. Omega-3 Index (O3i) was assessed using a dried blood spot sample in a subset of these athletes (n=228). Only 6% (n = 93) of athletes achieved the Academy of Nutrition & Dietetics’ recommendation to consume 500 mg of the ω-3 FA’s docosahexaenoic acid (DHA) and eicosapentaenoic acid (EPA) per day. Use of ω-3 FA supplements was reported by 15% (n = 229) of participants. O3i was 4.33 ± 0.81%, with zero participants meeting the O3i benchmark of 8% associated with the lowest risk of cardiovascular disease. Every additional weekly serving of fish or seafood was associated with an absolute O3i increase of 0.27%. Overall, sub-optimal ω-3 FA status was observed among a large, geographically diverse group of male and female collegiate athletes. These findings may inform interventions aimed at improving ω-3 FA status of collegiate athletes. Further research on athlete-specific ω-3 FA requirements is needed.

## Introduction

Omega-3 polyunsaturated fatty acids (ω-3 FA), namely long-chain eicosapentaenoic acid (EPA) and docosahexaenoic acid (DHA), serve as structural components within phospholipid cell membranes. These ω-3 FA have also been shown to play important physiological roles among the cardiovascular,(1–6) nervous,(7–13) and skeletal muscle systems(14–18), and in the body’s inflammatory response.(19–26) In athletes, ω-3 FA have been associated with the management of exercise-induced oxidative stress,(19,20,23–25) delayed onset muscle soreness,(21,22,25,26) oxygen efficiency during aerobic exercise,(2) anaerobic endurance capacity,(3) and skeletal muscle health.(14–18) The potential neuroprotective role of DHA as related to concussion and traumatic brain injury (TBI) has also been investigated.(8–13)

As essential fats, EPA and DHA must be obtained exogenously because the human body has limited ability to synthesize these ω-3 FA from precursor ω-3 FA alpha-linolenic acid (ALA).(27) Fish are the richest sources of ω-3 FA, but there is wide variation in the EPA and DHA content of these foods (Table 1). (28–32) Also of note, some commonly consumed sources like tuna and shellfish contain relatively smaller amounts of EPA and DHA and frequent consumption risks exposure to the effects of mercury. (28–32)

**Table 1.** Content of Omega-3 Fatty Acids in Common Food Sources.

| **FOOD (100g unless otherwise stated)\*\*** | **EPA+DHA (g)** | **EPA (g)** | **DHA (g)** | **ALA (g)** |
| --- | --- | --- | --- | --- |
| Cod Liver Oil (1 Tablespoon) | 2.43 | 0.938 | 1.492 | 0.042 |
| Salmon | 2.147 | 0.69 | 1.457 | 0.113 |
| Herring | 2.125 | 1.242 | 0.883 | 0.073 |
| Whitefish | 1.612 | 0.406 | 1.206 | 0.235 |
| Sardines | 1 | 0.4 | 0.6 | 0.5 |
| Bluefish | 0.988 | 0.323 | 0.665 | 0 |
| Trout | 0.936 | 0.259 | 0.677 | 0.199 |
| Swordfish | 0.819 | 0.138 | 0.681 | 0.238 |
| Bass | 0.763 | 0.305 | 0.458 | 0.142 |
| Whiting | 0.518 | 0.283 | 0.235 | 0.013 |
| Flounder | 0.501 | 0.243 | 0.258 | 0.016 |
| Mussels | 0.5 | 0.2 | 0.3 | 0 |
| Lobster | 0.48 | 0.341 | 0.139 | 0.01 |
| Sea Trout | 0.476 | 0.211 | 0.265 | 0.005 |
| Crab | 0.474 | 0.243 | 0.231 | 0.021 |
| Halibut | 0.465 | 0.091 | 0.374 | 0.083 |
| Oysters | 0.44 | 0.229 | 0.211 | 0.063 |
| Mackerel | 0.401 | 0.174 | 0.227 | 0 |
| Snapper | 0.321 | 0.048 | 0.273 | 0 |
| Clams | 0.284 | 0.138 | 0.146 | 0.008 |
| Tuna | 0.279 | 0.047 | 0.232 | 0.015 |
| Cod | 0.276 | 0.103 | 0.173 | 0.003 |
| Haddock | 0.238 | 0.076 | 0.162 | 0.003 |
| Catfish | 0.237 | 0.1 | 0.137 | 0.096 |
| Scallops | 0.189 | 0.086 | 0.103 | 0 |
| Shrimp | 0.164 | 0.02 | 0.144 | 0.012 |
| Mahi Mahi | 0.139 | 0.026 | 0.113 | 0.006 |
| Tilapia | 0.135 | 0.005 | 0.13 | 0.045 |
| Canola Oil (1 Tablespoon) | 0 | 0 | 0 | 1.2 |
| Flax Seeds (1 Tablespoon) | 0 | 0 | 0 | 2.2 |
| Chia Seeds (1 Tablespoon) | 0 | 0 | 0 | 2.673 |
| Flaxseed Oil (1 Tablespoon) | 0 | 0 | 0 | 8.5 |
| Walnuts | 0 | 0 | 0 | 3.306 |

\*Sorted from highest to lowest content of EPA+DHA
\*\*100 g is approximately equivalent to 3.5 ounces
\*\*\* ω-3 FA values using both USDA Addendum A: EPA and DHA Content of Fish Species and Bowes and Church's Food Values of Portions Commonly Used as reference.

While there is currently no consensus for ω-3 FA dietary recommendations (Table 2),(33–36) low ω-3 FA intake appears to be prevalent within the general population of North America, primarily attributed to the limited number of food sources, which includes a short list of fish and seafood, and infrequent consumption of these ω-3 FA rich foods.(28,29,37,38) Reports of athletes’ dietary ω-3 FA intake are minimal to date, but Wilson and Madrigal(39) observed intakes of EPA and DHA below 100 mg daily in a group of 58 National Collegiate Athletics Association (NCAA) Division I collegiate athletes. While no athlete-specific guidelines have been established, this is significantly less than the recommendation from the Academy of Nutrition & Dietetics to consume two fish servings weekly, providing a daily average of 500 mg EPA + DHA.(33) Little information is available about athletes’ habitual use of ω-3 FA supplements.

**Table 2.** Omega-3 Fatty Acid Dietary Recommendations for the General Public.

|  | Year of Publication | Population | Dietary Recommendation |
| --- | --- | --- | --- |
| Academy of Nutrition & Dietetics | 2014 | General Public (adults) | 500 mg EPA+DHA/ day<sup>34</sup> |
| American Heart Association | 2002 | General Public | At least 2 fish servings weekly (3.5 oz per serving)<sup>35</sup> |
| World Health Organization | 2003 | General public (adults) | 1-2% of energy/ day<sup>36</sup> |
| National Academy of Medicine (formerly Institute of Medicine) | 2005 | Adult men | 1.6 g/ day of ALA, of which ~10% EPA+DHA<sup>37</sup> |
|  |  | Adult women | 1.1 g/ day of ALA, of which ~10% EPA+DHA<sup>37</sup> |

In addition to ω-3 FA intake, ω-3 FA status may be evaluated using the Omega-3 Index (O3i), which reflects the sum of EPA and DHA in erythrocyte membranes as a percentage of total erythrocyte fatty acids.(4) Compared to other methods, O3i requires a minimum amount of blood (i.e., finger stick blood sample), has a low biological variability,(40) is less affected by acute feedings to better reflect long-term ω-3 FA intake,(41) and has been shown to correspond with ω-3 FA concentrations in the heart, brain, and a variety of other tissues.(42,43) An O3i <4% has been associated with the highest risk for the development of cardiovascular disease; whereas, 4-8% is considered moderate risk and ≥8% is the lowest risk.(4–6) Recently, an average O3i of 4.4% was observed among collegiate football athletes at four U.S. universities,(44) however a large scale assessment of O3i including non-football athletes has not been described in the peer reviewed published literature, to our knowledge.

Prior to 2019, the NCAA considered ω-3 FA supplements to be “impermissible”, which prevented athletic departments from purchasing such supplements for student-athletes. However, recently amended NCAA legislation reclassified ω-3 FA supplements, permitting athletic departments to provide them to student-athletes.(45) As a result of this rule change, interest in and availability of ω-3 FA supplements has risen. In order to better inform recommendations and ultimately nutrition interventions, a better understanding of athletes’ ω-3 FA status is needed. Thus, the purpose of this study was to assess the ω-3 FA intake and O3i of male and female NCAA Division I collegiate student-athletes who participate in a variety of sports.

## Methods

### Study Design

A multi-site, cross-sectional study was designed to assess the ω-3 FA dietary intake, ω-3 FA supplement use, and O3i of collegiate student-athletes. These assessments were carried out during the 2018-2019 academic year.

### Participants

Student-athletes from nine NCAA Division I institutions were invited to participate in the study. In order to achieve geographical diversity, institutions were dispersed throughout the U.S. (California, Georgia, Illinois, Nebraska, Oregon, Pennsylvania, Texas, Utah and Virginia). All nine institutions were classified as Power 5 programs. Male and female student-athletes who were over the age of 18 years and on a current roster for any NCAA Division I sport at one of the participating institutions were eligible to participate.

### Omega-3 Dietary Assessment

A 26-item food frequency questionnaire (FFQ) validated to assess ω-3 FA dietary intake(39,46) was administered to participants. The FFQ was modified to include demographic characteristics of participants (sex, age, academic year, and sport) and ω-3 FA supplement use. Within the FFQ, participants reported the frequency of consumption and average portion size for an extensive list of ω-3 FA food sources including fish, shellfish, walnuts, canola oil, flaxseed, flaxseed oil, and cod liver oil. For participants who indicated that they consumed ω-3 FA supplements, information about brand, form, dosage, and frequency taken was requested.

The FFQ results were compiled and analyzed using methodology outlined by Sublette et al.(46) Previously published databases(30–32) were used as a reference for ω-3 FA content of foods consumed based on source and portion size reported.

### Blood Fatty Acid Analysis

Following completion of the dietary assessment portion of the study, participants were offered the opportunity to volunteer for a second portion of the study: analysis of blood fatty acids. For the collection, a single drop of whole blood was sampled and applied to a blood spot card pre-treated with an antioxidant cocktail. Samples were shipped to a central laboratory (OmegaQuant, Sioux Falls, SD) for a full fatty acid analysis in addition to the calculation of the O3i using gas chromatography. This methodology is described in detail by Harris and Polreis.(47) The fatty acid analysis also included EPA, DHA and ALA.

### Statistical Analysis

Data were analyzed using IBM Statistical Package for the Social Sciences (SPSS) version 26. Descriptive statistics are expressed as means and standard deviations for continuous data, and frequencies and percentages for categorical data. Data were tested for normality using the Shapiro-Wilk test. Differences in outcomes between demographic groups were calculated using analysis of variance (ANOVA) or chi-square tests. Relationships between diet and blood variables were analyzed using Pearson’s correlations. Multiple regression analysis was used to assess the effects of diet on O3i after adjusting for demographic covariates. Significance was set at a level of p<0.05.

### Ethical Considerations

This study was approved by the Institutional Review Board of Virginia Tech (IRB# 18-606) and respective institutional research review committees. Consent for the dietary assessment portion of the study was inferred based on voluntary completion. Written and informed consent was provided by participants before starting the blood fatty acid portion of the study.

## Results

In all, 1528 participants (51% males) completed the dietary assessment portion of the study, from which 298 (55% males) completed the blood analysis portion. Participants represented 14 different male sports and 16 different female sports from nine institutions. Descriptive characteristics of participants are shown in Table 3. There were no differences in demographics between subject cohorts completing the dietary assessment and blood analysis portions of the study except that the blood cohort did not include ice hockey, ski, soccer, swimming & diving, and volleyball (male sports) and equestrian, field hockey, golf (female sports), and the Pennsylvania institution did not participate in the blood analysis (Table 3).

**Table 3.** Descriptive Characteristics of Participants.

|  | Dietary Assessment | Blood Fatty Acid Analysis | Differences Test Statistic (p-value) |
| --- | --- | --- | --- |
| **n** | 1,528 | 298 |  |
| **Sex (Male/Female)** | 780/748 | 163/115 | $\chi^2=1.318/$ (p=0.251) |
| **Age (years; mean ± sd)** | 19.9 ± 1.4 | 20.0 ± 1.3 | F=1.610/ (p=0.646) |
| **Academic year n (%)** | *Freshman: 442 (28.9%)*<br>*Sophomore: 373 (24.4%)*<br>*Junior: 377 (24.7%)*<br>*Senior: 270 (17.7%)*<br>*5<sup>th</sup> year or Graduate: 63 (4.1%)* | *Freshman: 88 (29.5%)*<br>*Sophomore: 73 (24.4%)*<br>*Junior: 70 (23.5%)*<br>*Senior: 58 (19.4%)*<br>*5<sup>th</sup> year or Graduate: 9 (3.0%)* | $\chi^2=18.50/$ (p=0.470) |
| **Sport n (%)** | *Football: 303 (19.8%)*<br>*<sup>a</sup> Non-football Male Sport: 477 (31.2%)*<br>*<sup>b</sup> Female Sport: 748 (49.0%)* | *Football: 81 (27.2%)*<br>*<sup>c</sup> Non-football Male Sport: 82 (27.5%)*<br>*<sup>d</sup> Female Sport: 115 (38.5%)* | $\chi^2=4.779/$ (p=.1912) |
| **Region n (5)** | *California: 106 (6.9%)*<br>*Georgia: 158 (10.3%)*<br>*Illinois: 77 (5.0%)*<br>*Nebraska: 211 (13.8%)*<br>*Oregon: 111 (7.3%)*<br>*Pennsylvania: 61 (4.0%)*<br>*Texas: 336 (22.0%)*<br>*Utah: 102 (6.7%)*<br>*Virginia: 365 (23.9%)* | *California: 28 (28.6%)*<br>*Georgia: 33 (11.1%)*<br>*Illinois: 29 (9.7%)*<br>*Nebraska: 45 (15.1%)*<br>*Oregon: 39 (13.1%)*<br>*Pennsylvania: 0 (0.0%)*<br>*Texas: 40 (13.4%)*<br>*Utah: 42 (14.1%)*<br>*Virginia: 43 (14.4%)* | $\chi^2=7.003/$ (p=.0991) |

<sup>a</sup> Baseball, Basketball, Cross Country, Golf, Gymnastics, Ice Hockey, Ski, Soccer, Swimming & Diving, Tennis, Track & Field, Volleyball, Wrestling
<sup>b</sup> Basketball, Bowling, Cross Country, Cheerleading, Equestrian, Fencing, Field Hockey, Golf, Gymnastics, Lacrosse, Rifle, Rowing, Soccer, Softball, Swimming & Diving, Track & Field
<sup>c</sup> Baseball, Basketball, Cross Country, Golf, Gymnastics, Tennis, Track & Field, Wrestling
<sup>d</sup> Basketball, Cross Country, Cheerleading Fencing, Gymnastics, Lacrosse, Rifle, Rowing, Soccer, Softball, Swimming & Diving, Track & Field

### Diet

A total of 659 participants (45%) reported consuming no fish in the last 6 months, the most significant source of DHA and EPA. A total of reported per The AHA’s recommendation of consuming at least two or more fish servings weekly was met by 601 participants (39%). (34) Comparatively when considering all DHA and EPA sources, fish and/or seafood was consumed by 1345 participants (88%) at least once during the previous 6 month timeframe (Figure 1). Salmon and shrimp were the only EPA and DHA sources reported to be consumed by more than 50% of participants (Figure 2). ALA consumption included canola oil (85%), walnuts (53.9%), chia (43.6%), flax or flax oil (34.9%), and cod liver oil (3.3%). Use of ω-3 FA supplements was reported by 229 participants (15%). Of supplement-users, 153 (67%) purchased the supplement on their own, while 76 (33%) received supplements via their respective athletic program. Most participants provided no response to brand, type, and dose of ω-3 FA supplements consumed.

**Figure 1.**
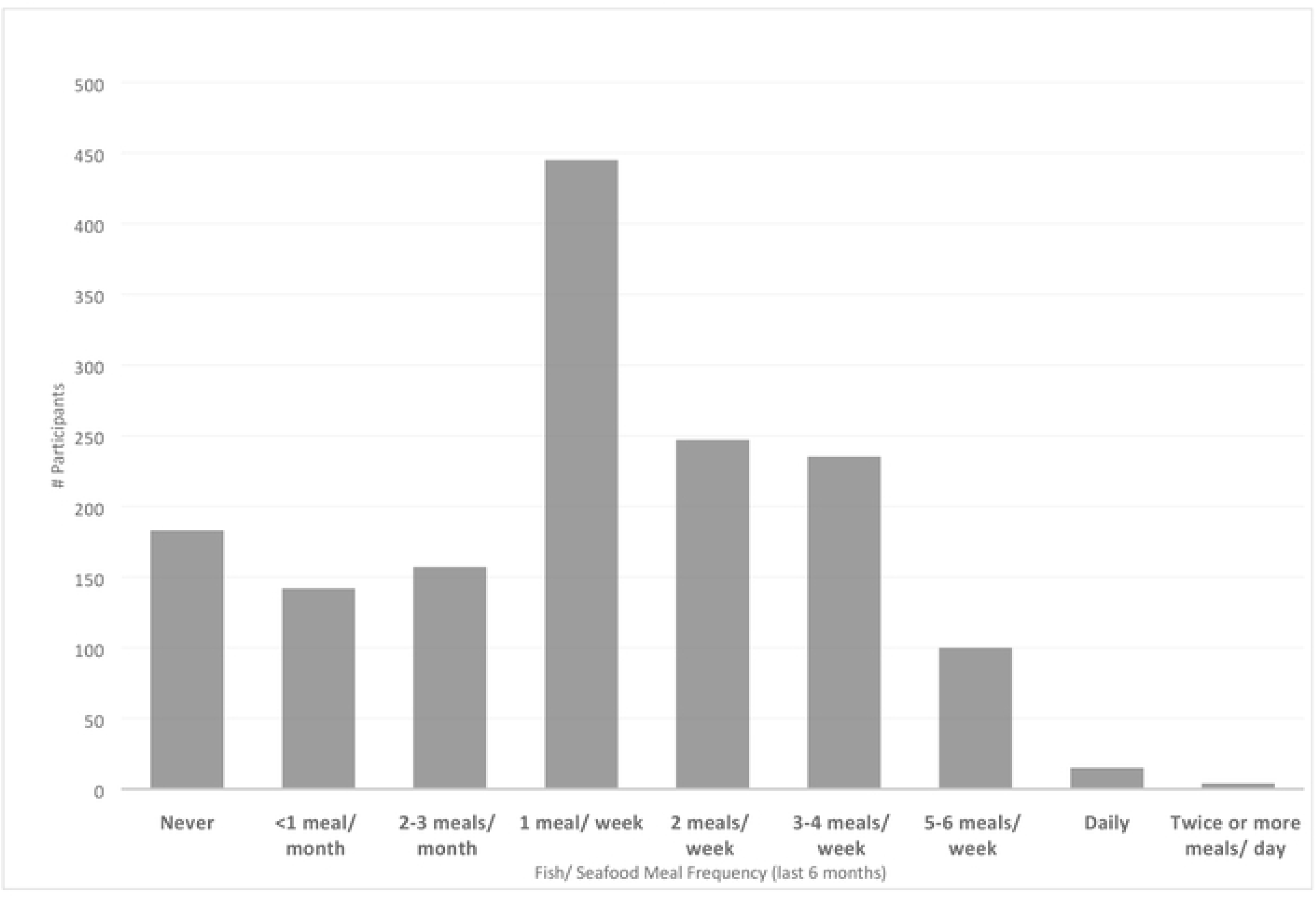
Frequency of Fish and Seafood Consumption During the Previous 6 Months (n = 1528)

**Figure 2.**
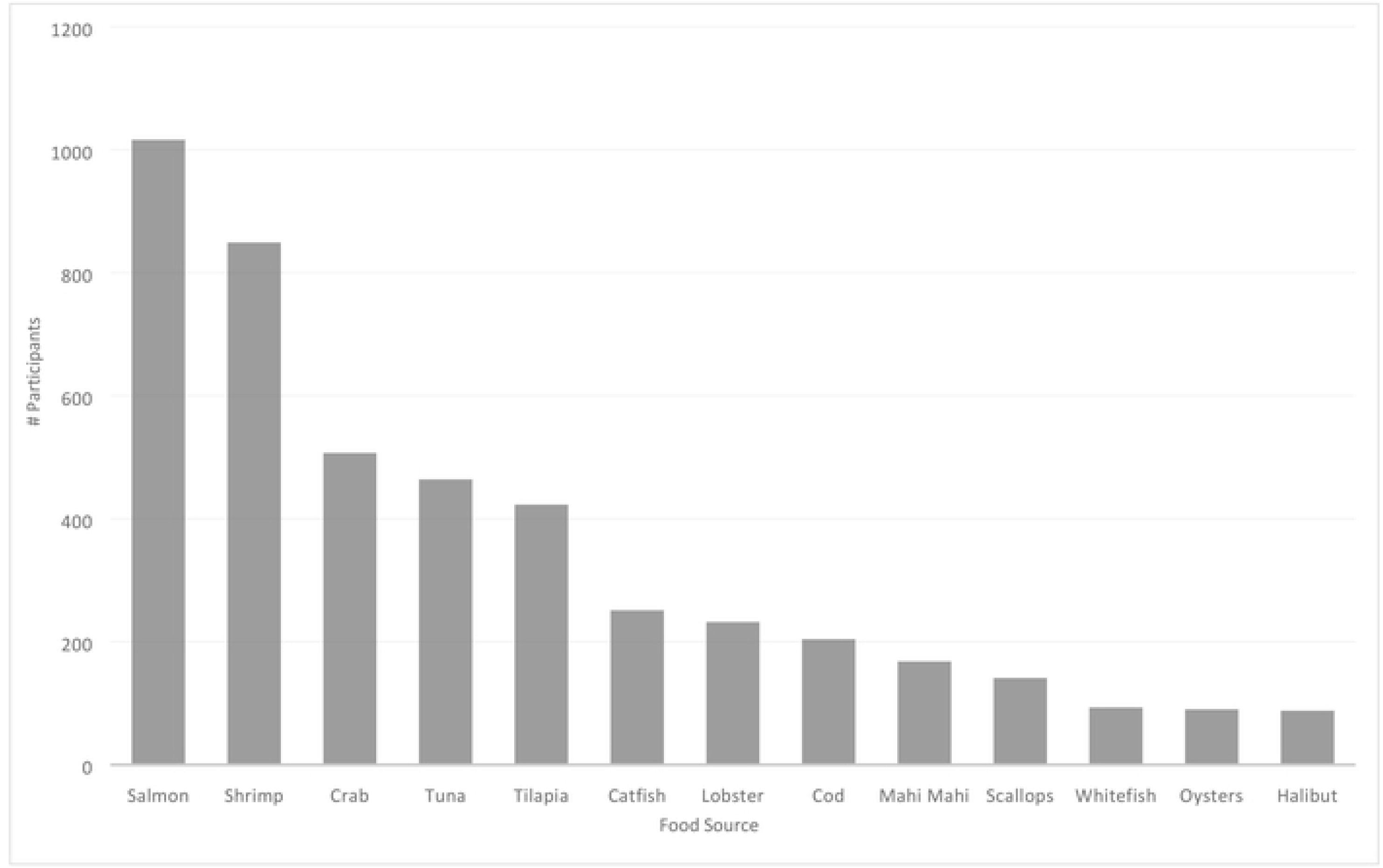
Sources of Fish and Seafood Consumption During the Previous 6 Months (n = 1528)

In comparison to previously reported dietary intake recommendations (Table 1), only 6% of participants consumed at least 500 mg EPA + DHA/day as advised by the Academy of Nutrition and Dietetics(33) and 4% met the National Academy of Medicine’s (formerly Institute of Medicine) recommendation of 1.6 g ALA (men) or 1.1 g ALA (women) with 10% coming from EPA + DHA.(36) Dietary consumption of EPA, DHA, EPA + DHA, and ALA are shown in Table 4.

**Table 4.** Dietary Consumption of Omega-3 Fatty Acids (n=1528)

|  | Total Daily Intake (mg) | Sex |  | p-value | Sport |  | p-value |
| --- | --- | --- | --- | --- | --- | --- | --- |
|  |  | Male<br>n= 780 | Female<br>n= 748 |  | Football<br>n= 303 | Non-football<br>n= 477 |  |
| EPA | 46.8 +/- 86.9 | 53.4 | 40.4 | .0042** | 57.3 | 43.6 | 0.0198* |
| DHA | 94.8 +/- 164.9 | 106.4 | 83.9 | .0091** | 111.8 | 89.7 | 0.0467* |
| ALA | 571.8 +/- 1151.5 | 530.4 | 626.6 | .0281* | 622.7 | 587.8 | 0.6621 |
| EPA + DHA | 141.7 +/- 250.6 | 159.8 | 124.3 | .0068** | 169.1 | 133.4 | 0.0342* |

### Blood

Result of blood EPA, DHA, ALA and O3i analyses are shown in Table 5. O3i ranged from 2.25 to 7.23% (Figure 3), with 114 (38%) in the high risk category, 184 (62%) in the moderate risk category, and 0 (0%) in the low risk category. There were no significant differences in blood measures based on sex (Figure 4), sport (Figure 5), location, age, or academic year.

**Table 5.** Blood Fatty Acid Analysis Results (n= 298)

|  | Blood Fatty Acids (%) | Sex |  | p-value | Sport |  | p-value |
| --- | --- | --- | --- | --- | --- | --- | --- |
|  |  | Male | Female |  | Football | Non-football |  |
| EPA | 0.45 ± 0.19 | 53.4 | 40.4 | p= 0.704 | 57.3 | 43.6 | 0.738 |
| DHA | 2.19 ± 0.59 | 106.4 | 83.9 | p= 0.699 | 111.8 | 89.7 | 0.718 |
| ALA | 0.49 ± 0.19 | 530.4 | 626.6 | p= 0.588 | 622.7 | 587.8 | 0.820 |
| Omega-3 Index (O3i) | 4.3 ± 0.81 | 4.3 | 4.4 | p= 0.905 | 4.4 | 4.4 | 0.942 |

**Figure 3.**
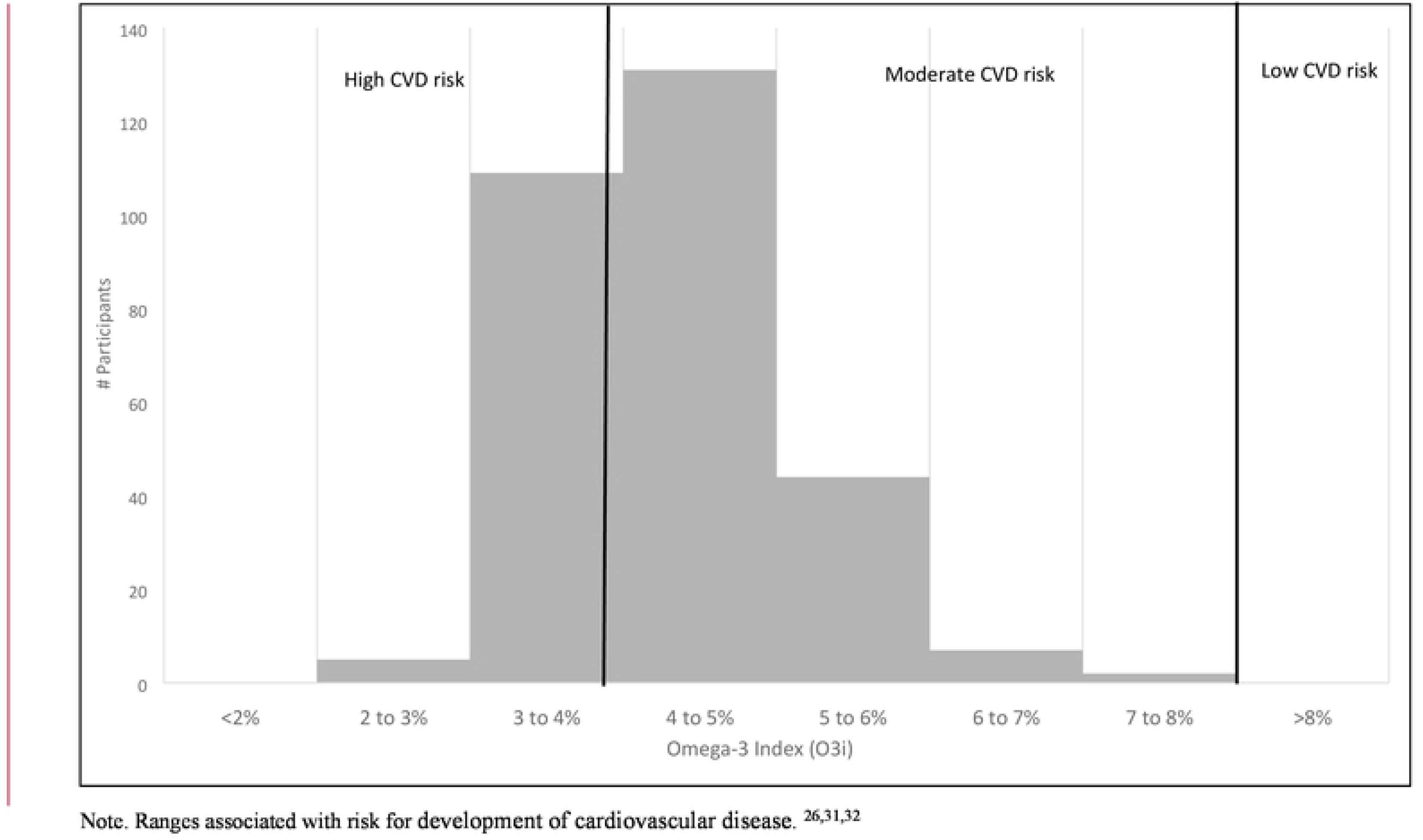
Distribution of Omega-3 Index Results (n=298)

**Figure 4.**
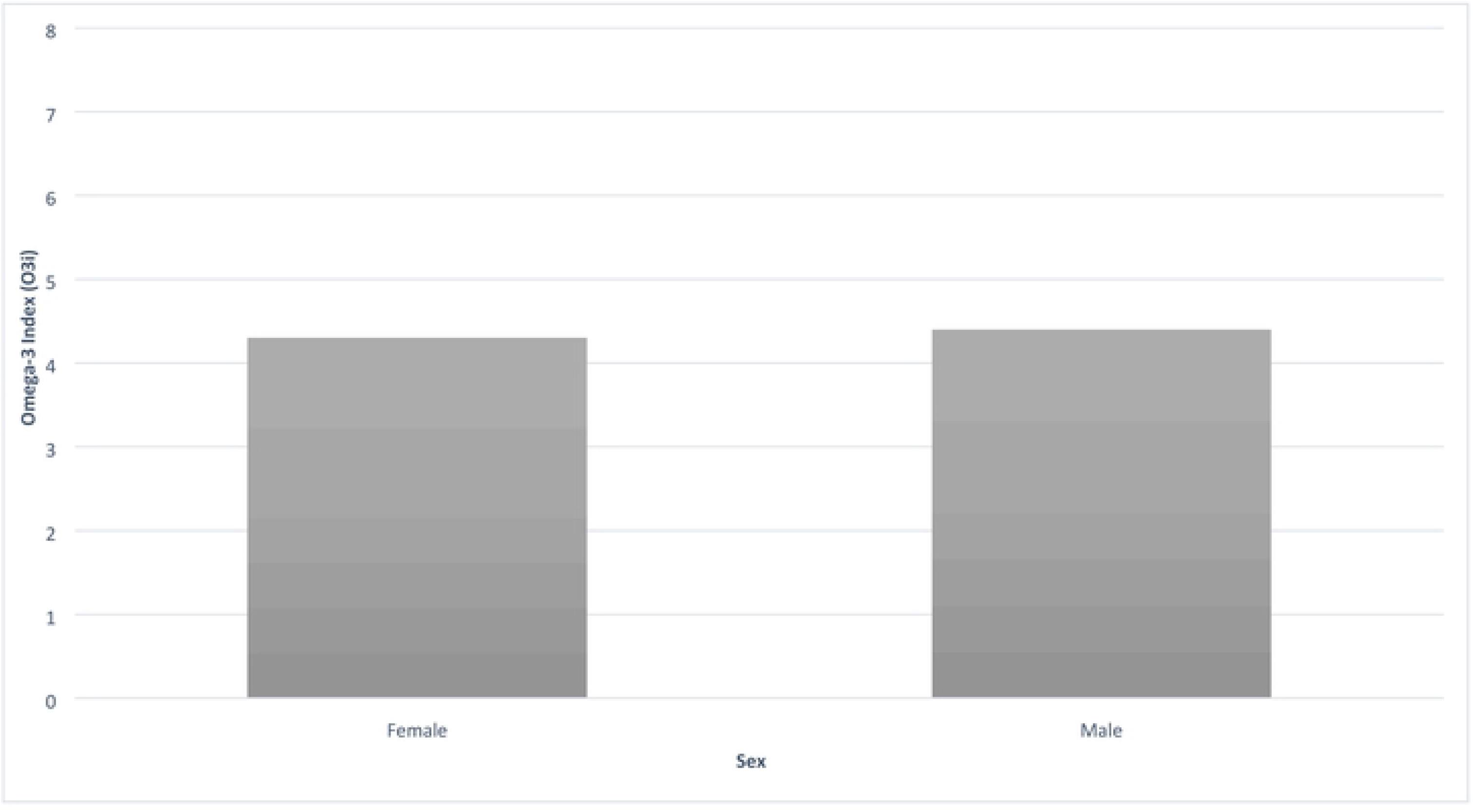
Average Omega-3 Index between Sexes.

**Figure 5.**
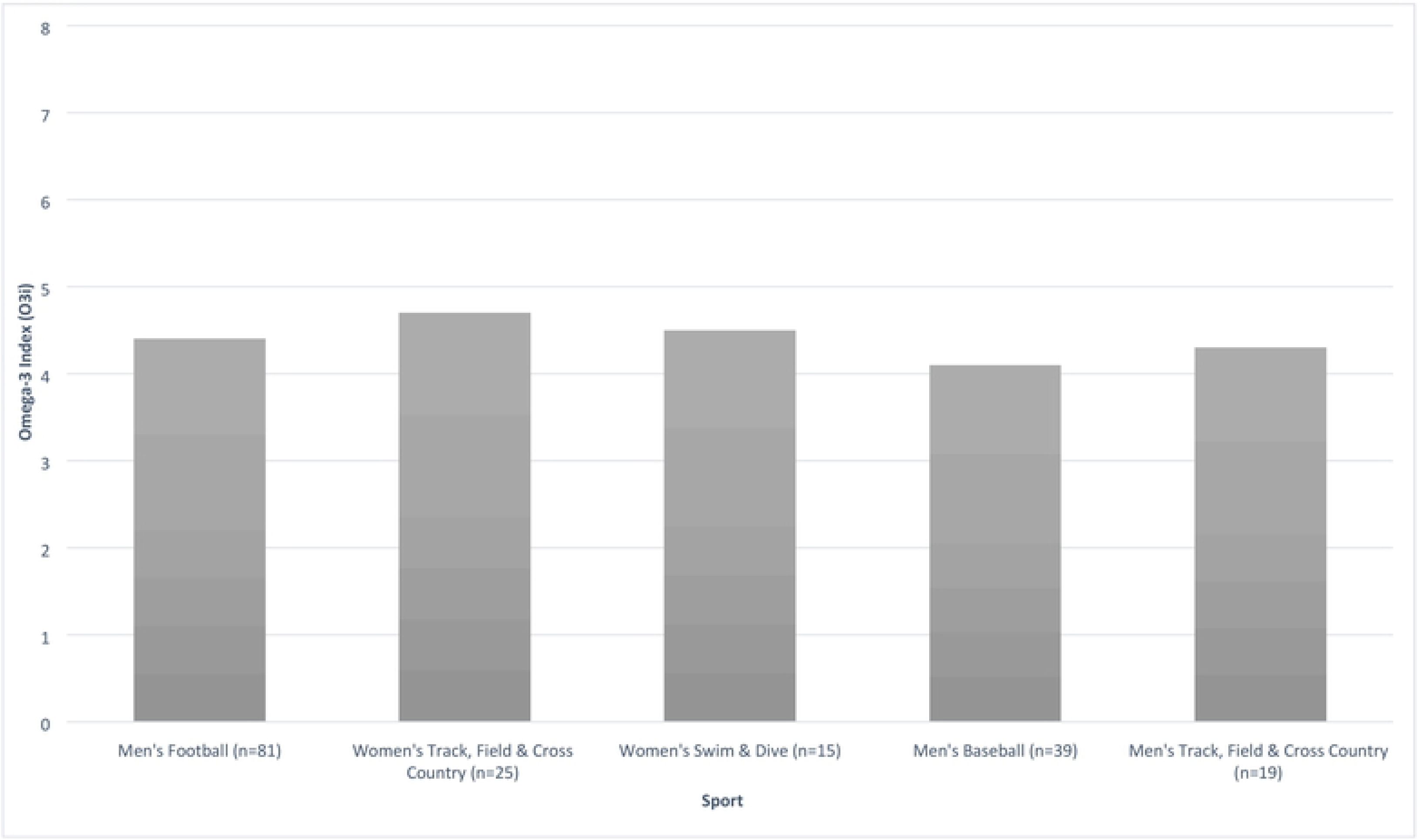
Average Omega-3 Index between 5Highest Participating Sports.

**Figure 6.**
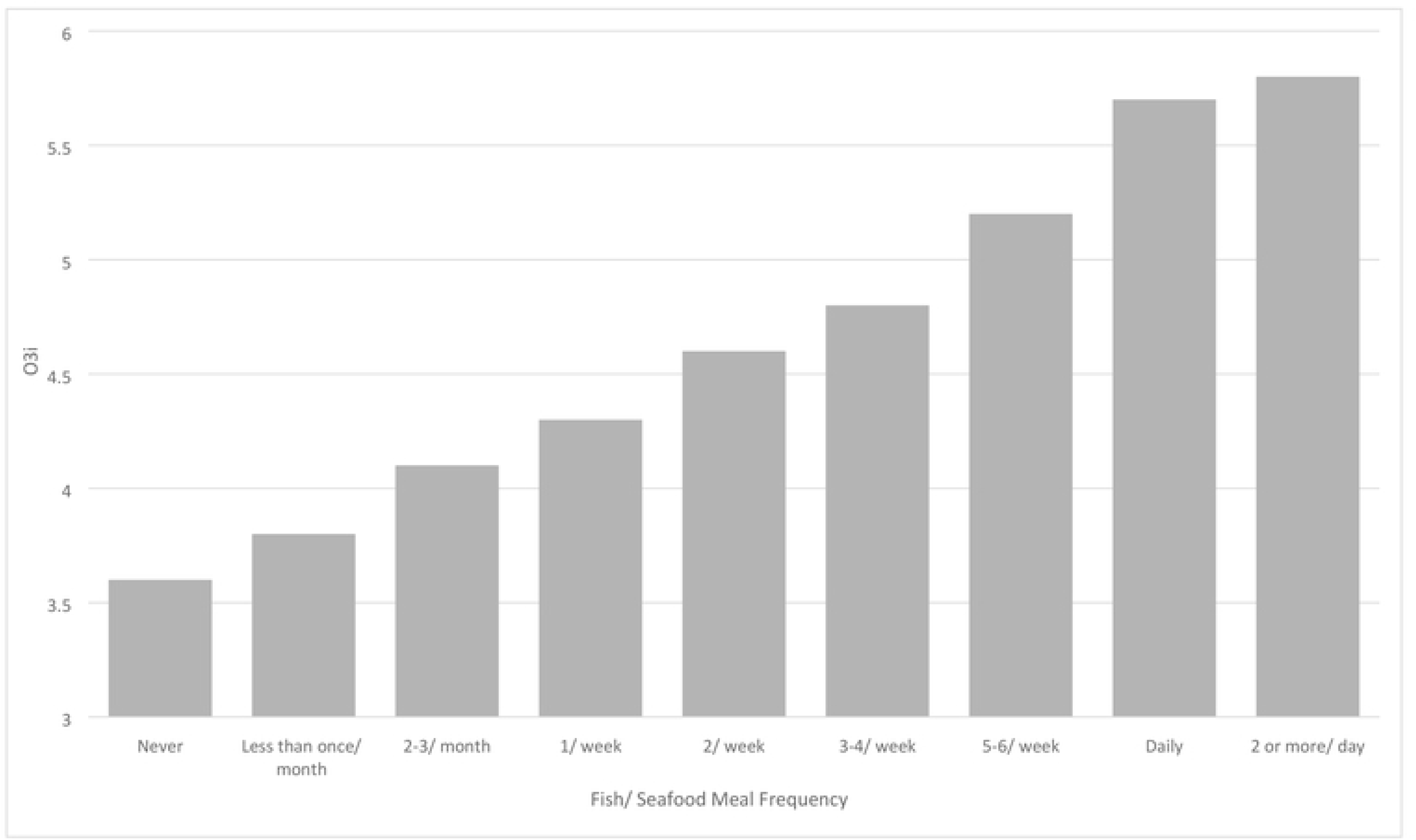
Relationship between Fish and Seafood Meal Frequency and Omega-3 Index (n = 298)

### Relationship Between Diet and Blood Measures

Dietary intake of both EPA and DHA were positively correlated with blood EPA, DHA, and O3i (Table 6). Dietary ALA intake had no correlation with blood levels of EPA, DHA, ALA or O3i (Table 6).

**Table 6.** Diet and Blood Fatty Acid Correlations Table.

|  | Diet EPA | Diet DHA | Diet EPA + DHA | Diet ALA | Diet Total ω-3 | Blood EPA | Blood DHA | Blood ALA | Blood ω-3 Index |
| --- | --- | --- | --- | --- | --- | --- | --- | --- | --- |
| **Diet EPA** | 1 |  |  |  |  |  |  |  |  |
| **Diet DHA** | .977 ** | 1 |  |  |  |  |  |  |  |
| **Diet EPA + DHA** | 0.990 ** | 0.997 ** | 1 |  |  |  |  |  |  |
| **Diet ALA** | 0.134 | 0.154 | 0.148 | 1 |  |  |  |  |  |
| **Diet Total ω-3** | 0.332 * | 0.552 * | 0.347 * | 0.979 ** | 1 |  |  |  |  |
| **Blood EPA** | 0.342 * | 0.334 * | 0.338 * | 0.296 | 0.339 * | 1 |  |  |  |
| **Blood DHA** | 0.397<br>* | 0.404<br>* | 0.403<br>* | 0.214 | .273 | .402 * | 1 |  |  |
| **Blood ALA** | 0.072 | 0.080 | 0.078 | 0.090 | 0.098 | 0.072 | -0.122 | 1 |  |
| **Blood ω-3 Index** | 0.437<br>* | 0.441<br>* | 0.442<br>* | 0.271 | 0.332<br>* | 0.648<br>** | 0.958<br>** | -0.079 | 1 |

**Note: \*p<0.05, \*\*p<0.01**

After controlling for location, sex, age, class year and sport (football vs. non-football), frequency of seafood consumption was a significant predictor of O3i (R^2^=.3701, p<0.01). Each additional serving of seafood was associated with a O3i increase of 0.27% (Figure 5). Participants who reported taking ω-3 FA supplements had significantly higher O3i compared with those not taking supplements (4.7 vs. 3.7%, respectively; p<0.05). Participants who met the Academy of Nutrition and Dietetics’ recommendation of 500 mg EPA+DHA per day had a higher O3i on average compared to those who consumed less than the 500 mg EPA+DHA recommendation (5.4% vs. 4.3%, p<0.05).

## Discussion

The primary goal of this study was to describe the ω-3 FA status of collegiate athletes in the U.S. Our findings indicate that collegiate athletes are not meeting dietary recommendations for ω-3 FA and have sub-optimal O3i as compared to currently proposed cardiovascular benchmarks. To our knowledge, this is the first large scale assessment of ω-3 FA status of male and female collegiate athletes from a variety of sports.

While the majority of collegiate athletes participating in the present study did not meet current dietary ω-3 FA recommendations (Table 1), similar to previous observations(44,46) it is important to note that these guidelines are not specific to athletes. Further research is needed to establish athlete-specific recommendations, especially taking into consideration the physiological implications of advanced levels of training on metabolism and the inflammatory response. (48–50) For example, lower average O3i was observed among non-elite runners with greater training mileage compared to those with lesser running mileage.(48)

Given the pattern of low ω-3 FA intake observed in collegiate athletes,(39,44) clinicians should consider nutritional interventions aimed at improving ω-3 FA status. One strategy could be increasing consumption of fish and seafood, the richest sources of EPA + DHA, as nearly half of participants reported no fish consumption in the last 6 months. In recent years, the NCAA has seen significant changes in terms of the feeding opportunities available for athletes as a result of the deregulation of meal restrictions on Division I collegiate student-athletes in 2014.(51) Based on our findings, inclusion of ω-3 FA-rich sources in provided meals is advisable. Capitalizing on popular fish and seafood sources (salmon, shrimp, crab, tuna, and tilapia were consumed the most in the current study) may be beneficial. Those involved in nutrition programming and meal planning should also recognize that plant-based sources of ω-3 FA are rich in ALA rather than EPA + DHA and that the conversion of ALA to EPA + DHA is minimal. (27) The observed lack of correlation between dietary ALA and blood measures of EPA, DHA and O3i, is also consistent with previous findings (39,46)

No participant in the current study, including those who consumed fish or seafood twice or more per week, had an O3i of 8%, the level associated with lowest cardiovascular disease risk.(4–6) Thus, achieving optimal ω-3 FA status through diet alone may be difficult and it is plausible that athletes may actually have higher needs than the general population. The use of ω-3 FA supplements is another strategy for improving ω-3 FA status, and has been discussed as a potentially helpful nutritional tool for athletes.(52) A small percentage of participants reported ω-3 FA supplement use but almost none were able to provide information about brand, form, dosage, and frequency of supplements used. The recent NCAA guidelines changes(45) present an opportunity to more readily provide ω-3 FA when appropriate for student-athletes, and to do so in a safe, controlled, and monitored fashion.

The sub-optimal O3i observed for in our study (4.3%) was similar to previous observations,(39,44,53,54) and did not differ based on sex or sport. While further research is needed to investigate potential differences in needs between athletes of different sex and sport, these we observed collegiate athletes collectively have low ω-3 FA status. Higher consumption of EPA+DHA observed in males and football participants compared to their counterparts did not translate to higher O3i values. This might suggest external factors such as higher average body mass, higher caloric needs and availability of athletic department nutrition resources drove the observed increases in EPA+DHA intake and was not significant enough to impact blood status. To our knowledge, no U.S.-based athletes have been documented in the peer reviewed literature as having O3i greater than 8%,(39,44) the proposed benchmark for optimal cardiovascular health.(4–6) Given the increasing risk of cardiovascular disease reported among athletes,(55) a focus on improved O3i is warranted. Although O3i is positively correlated with ω-3 FA concentration of a variety of tissues, and ω-3 FA status is associated with a number of health and performance factors for athletes,(2,3,8–13,15–24,26) target O3i for non-cardiovascular conditions is not well-established. Continuing research is needed to investigate the impact of O3i on athlete health and performance measures.

### Strengths & Limitations

Collaboration with a diverse group of Power 5 institutions enabled us to study a large sample of athletes from nearly every NCAA sport with varying dietary habits and available resources. Further, given the timing of the NCAA legislation changes in relation to the timeline of our assessment, this investigation also serves as a baseline for ω-3 FA intake and ω-3 FA supplement use among collegiate athletes. Finally, our results parallel those of others who have observed a positive correlation between dietary EPA and DHA intake and O3i.(56–58) This suggests that the FFQ we used(39,46) was a reliable measure of ω-3 FA intake. This FFQ provides a cost-effective method for assessing ω-3 FA status in clinical situations where blood assessment may not be practically or financially warranted.

The study does have some limitations, however. For example, fish and seafood vary in nutritional content based on a number of factors, including variety consumed, location, and time of year. Our assessment did not account for this variation. Additionally, we did not collect data related to race/ethnicity, height, and body weight in effort to assure anonymity of participants, but this information may have been insightful in data analysis. Overall, the lack of universally accepted dietary recommendations and blood measure standards provided an additional obstacle in terms of interpreting our results, which should be a primary motive for future research.

### Conclusions

Prior to the change in NCAA legislation change related to ω-3 FA supplementation, we observed sub-optimal omega-3 status in NCAA Division I athletes based on both dietary and blood assessments. These results serve to inform future nutritional interventions aimed at improving ω-3 FA status among athletes. Results also provide a baseline in order to measure the impact of nutrition interventions created as a result of this legislation change.

## Acknowledgements

The authors would like to thank dietitians and sport scientists at the nine participating institutions who assisted in the data collection process. This study was funded, in part, by the Collegiate and Professional Sports Dietitians Association Research Award.

